# Predicting host-pathogen interactions using a proteome-scale language model

**DOI:** 10.64898/2026.05.29.728699

**Authors:** Cyril Malbranke, Cecilia Fruet, Anne-Florence Bitbol

## Abstract

ProteomeLM (Malbranke et al., 2025) is a proteome-scale language model trained on proteomes spanning the tree of life to reconstruct masked protein embeddings from proteome context within each species. Its attention coefficients capture protein-protein interactions without supervision. Here, we show that this capability extends to cross-species host-pathogen interactions (HPI) across ten human pathogen taxa spanning viruses and bacteria, and can be further improved with lightweight fine-tuning. We introduce **ProteomeLM-HPI**, a parameter-efficient adaptation via LoRA, trained on concatenated host-pathogen proteomes to reconstruct masked pathogen embeddings from host context. ProteomeLM-HPI involves two key design choices: *asymmetric masking* (pathogen-heavy masking) and *blocked self-attention*. Systematic ablations show that both choices contribute. To assess generalization, we introduce a strict cross-species benchmark enforcing pathogen-level hold-out and 40% sequence-identity filtering. On this benchmark, Proteome-HPI improves AUC on 9 out of 10 unseen pathogens.

## 1. Introduction

Knowing which proteins a pathogen deploys against its host, and which host proteins they target, is fundamental to understanding infection and pathogenesis. Mapping these HPI networks at the proteome scale would accelerate the identification of drug targets and vaccine antigens, yet large-scale experimental determination remains expensive and incomplete. Computational prediction is therefore essential, but HPI prediction is more challenging than intraspecies PPI prediction.

Host and pathogen proteins evolve under independent selective pressures and share no meaningful sequence homology. This precludes the use of amino-acid coevolution-based methods (including direct coupling analysis (DCA) (Weigt et al., 2009)), which require paired multiple sequence alignments, to predict cross-species interaction. Structure-based approaches such as AlphaFold-Multimer (Evans et al., 2021) can resolve individual interfaces with high accuracy but require a computationally intensive run for each candidate interaction pair, making proteome-wide screening impractical. Sequence-based supervised classifiers (Tsukiyama et al., 2021; Sledzieski et al., 2021; Singh et al., 2022; Liu et al., 2025) perform well on curated benchmarks, but often generalize poorly across taxa and can be confounded by sequence-identity leakage between training and test splits.

ProteomeLM (Malbranke et al., 2025) is a transformer-based language model trained on ∼32,000 complete proteomes spanning the tree of life. It takes as input the full set of proteins encoded by a genome, each represented by a pre-computed ESM-Cambrian (ESM-C) embedding (ESM Team et al., 2024), and reconstructs masked protein embeddings from their proteome context using a functional encoding based on orthologous groups (Kuznetsov et al., 2022) instead of positional encodings. A striking property emerges: ProteomeLM’s attention coefficients spontaneously encode PPI. For instance, a logistic regression on the 48 attention heads achieves an AUROC exceeding 0.9 for PPI recovery in *Escherichia coli*, without any interaction-level supervision (Malbranke et al., 2025). *Can the same masked reconstruction principle, applied to a concatenated host-pathogen proteome, predict cross-species HPI?*

Here we show that ProteomeLM’s attention heads already encode substantial cross-species HPI signal without any fine-tuning or interaction-level supervision (AUROC = 0.786 across ten pathogens). Furthermore, this signal can be amplified with fine-tuning. Specifically, we introduce **ProteomeLM-HPI**, which adapts ProteomeLM via LoRA (low-rank adaptation) (Hu et al., 2022) on paired host-pathogen proteomes using asymmetric masking and blocked pathogen self-attention. Through a systematic ablation across ten pathogens (six viruses and four bacteria), we show that these two design choices are beneficial and jointly optimal. As ProteomeLM-HPI processes the full host-pathogen proteome in one forward pass, it yields interaction predictions for all possible protein pairs simultaneously, which is more efficient than per-pair inference pipelines.

## 2. Methods: training ProteomeLM-HPI

### Input construction

Each training example pairs a host and a pathogen proteome. Proteins are represented as pre-computed ESM-C embeddings (dimension 1152).

### Masking strategy

ProteomeLM (Malbranke et al., 2025) applies uniform masking within a single-species proteome. For HPI fine-tuning, we apply distinct masking rates to host and pathogen proteins. When the host masking rate *m*_*H*_ is kept lower than the pathogen rate *m*_*P*_ ≥ *m*_*H*_, heavy pathogen masking forces the model to reconstruct pathogen proteins almost entirely from host proteome context, putting attention away from within-species dependencies toward cross-species ones. We evaluate four masking strategies: *symmetric* (*m*_*H*_ = *m*_*P*_ = 0.5), *asymmetric* (*m*_*H*_ = 0.2, *m*_*P*_ = 0.8), *total* (*m*_*H*_ = 0.2, *m*_*P*_ = 1.0), and *inverted* (*m*_*H*_ = 0.8, *m*_*P*_ = 0.2).

### Blocked pathogen self-attention

We zero all attention logits from pathogen tokens to other pathogen tokens at every layer, preventing pathogen proteins from attending to one another. This choice avoids a potential shortcut: without it, the model could reconstruct masked pathogen proteins by attending to unmasked pathogen proteins, bypassing any need to develop cross-species attention patterns.

### Dataset collection

We aggregated experimentally verified host-pathogen protein-protein interactions from PHISTO (Durmuş Tekir et al., 2013), IntAct (Del Toro et al., 2022), and HPIDB2 (Ammari et al., 2016), yielding 589 unique host-pathogen organism pairs, of which 59 are held out as a validation set to monitor training. To increase training diversity, we also developed and tested a data augmentation strategy where each seed pair is expanded to all organisms sharing a genus with either the host or the pathogen (NCBI Taxonomy), growing the pool to 32 484 host-pathogen proteome pairs. We evaluate performance on ten well-studied human pathogens for which large numbers of experimentally verified interactions are available: six viruses (SARS-CoV-2, HIV-1, Influenza A, EBV, HPV-16, HSV-1) and four bacteria (*Y. pestis, S. enterica* (salmonella), *C. trachomatis* (chlamydia), *M. tuberculosis*).

### Training protocol

We use ProteomeLM-S (36M parameters, 6 layers, 8 attention heads). We fine-tune it with LoRA (*r* = 32, *α* = 128, dropout 0.1) using the AdamW optimizer with a cosine schedule (lr 10^−5^, 500 warm-up steps, one H100 GPU). Ablation models are trained for 20 000 steps (1 hour); the final model for 80 000 steps (4 hours).

## 3. Results

### 3.1. ProteomeLM encodes cross-species HPI signal without supervision

Figure 1A shows the mean AUROC resulting from a logistic regression trained on the attention heads for predicting protein interaction across all ten pathogen datasets for each ablation configuration. To avoid inflating performance estimates due to leakage of sequence-similar proteins, we evaluate using grouped cross-validation: clusters are computed separately for pathogen and host proteins at 40% identity (MMseqs2 (Steinegger and Söding, 2018)), and each protein pair is assigned to its pathogen-cluster, host-cluster group, ensuring that no homologous protein appears simultaneously in training and evaluation sets. The base ProteomeLM achieves a mean AUROC of 0.786 across all ten pathogen datasets. This is a striking result: trained on single proteomes, and with no interaction-level supervision of any kind, ProteomeLM’s attention coefficients already encode substantial cross-species HPI signal. It confirms that proteome-scale pre-training over thousands of species induces representations that generalize.

**Figure 1.**
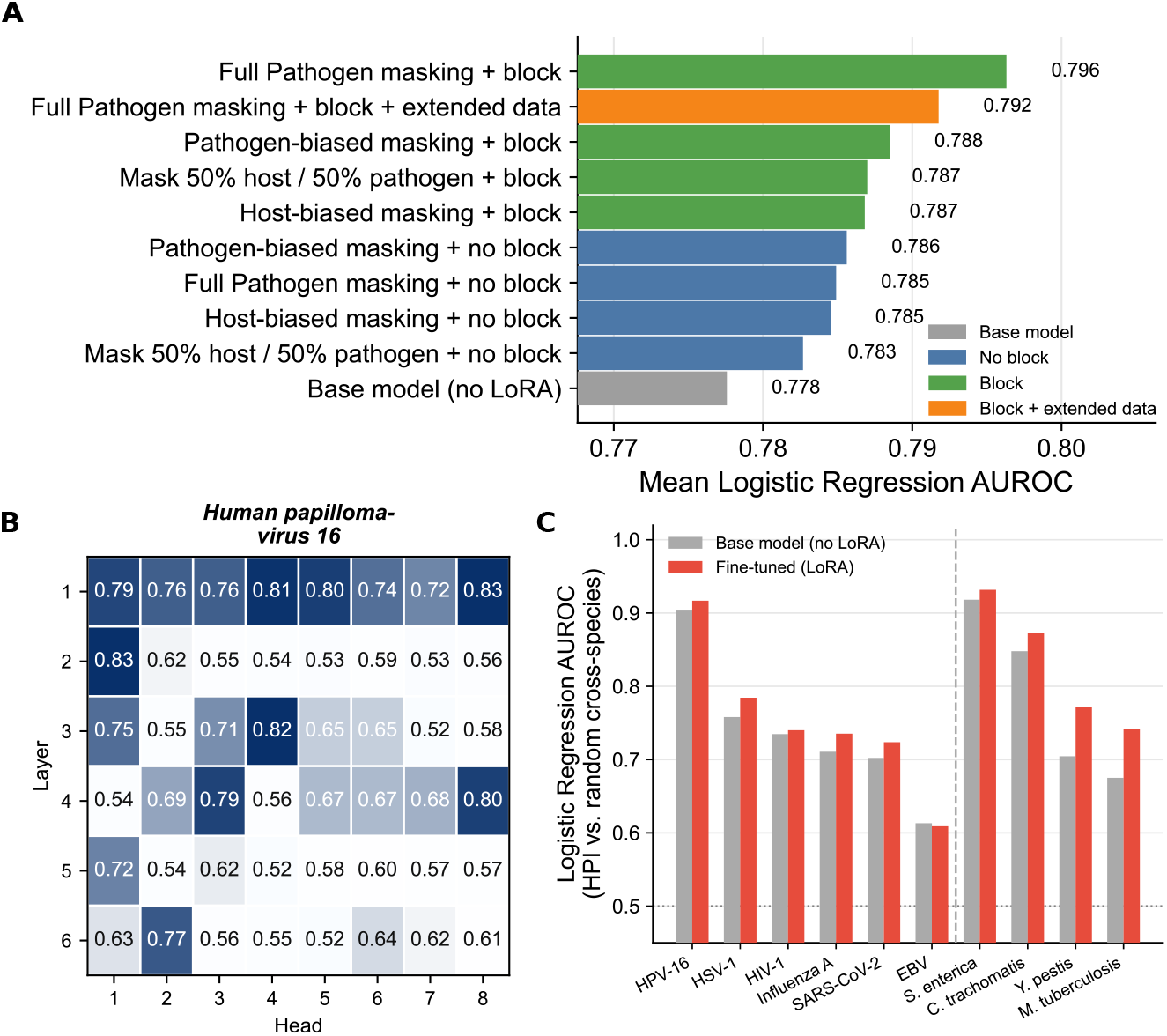
Main results and ablations for ProteomeLM-HPI. **A:** Mean logistic regression AUROC (HPI vs. random cross-species pairs) across ten pathogens for the base model and all fine-tuning ablations, evaluated via cross-validation (filtering by sequence similarity). **B:** Per-attention head AUROC (interacting vs. non-interacting pair) for human-HPV 16 interactions, using ProteomeLM-HPI (best setting from A). **C:** Per-dataset comparison between the best fine-tuned model and thebase model.

### 3.2. HPI signal is amplified by fine-tuning with asymmetric masking and blocked self-attention

All eight fine-tuned configurations (four masking strategies, with and without pathogen self-attention blocking) improve over the strong baseline obtained with ProteomeLM. Two independent axes of improvement are visible. First, blocking pathogen self-attention provides a consistent gain for every masking strategy. This confirms that preventing pathogen proteins from attending to one another is a beneficial bias: it forces the model to reconstruct masked pathogen proteins from host context rather than from within-pathogen signal, directly encouraging cross-species attention patterns. Second, among blocked models, increasing the pathogen masking fraction further raises performance: AUROC rises from 0.787 (symmetric, *m*_*P*_ = 0.5) to 0.796 (total, *m*_*P*_ = 1.0).

We also trained a model under the best setting using data augmentation obtained by expanding the seed pairs (590 pairs *→*32 484 pairs, see Methods). This did not improve performance, reaching a mean AUROC of 0.792 (versus 0.796 for the model trained on the smaller set). This suggests that the expansion introduces pairs with weaker or noisier cross-species signal, diluting the clean supervision carried by the curated seed interactions.

### 3.3. Fine-tuning generalizes across pathogen taxa

We trained ProteomeLM-S with total pathogen masking and blocked self-attention, and we called it ProteomeLM-HPI. Figure 1C shows per-dataset results comparing ProteomeLM-HPI against the ProteomeLM-S base model.

Fine-tuning improves HPI discrimination for nine out of ten pathogen datasets, with only a negligible degradation on EBV (0.613→ 0.609). This is compelling given the taxonomic breadth of the evaluation set: the model generalizes across diverse pathogens, including viruses and bacteria.

The largest gains are observed for *Y. pestis* (0.704 → 0.772), *M. tuberculosis* (0.675 → 0.742), and HSV-1 (0.758 → 0.784), while the highest absolute fine-tuned AUROCs are obtained for *S. enterica* (0.932), HPV-16 (0.917), and *C. trachomatis* (0.873). Fine-tuning tends to provide larger gains for pathogens with lower initial signal, suggesting that the inductive bias introduced by asymmetric masking and blocked self-attention is most useful when the base model’s cross-species representations are weakest.

### 3.4. Comparison with prior methods on an existing benchmark

Table 1 compares ProteomeLM-HPI against published methods on the benchmark of (Tsukiyama et al., 2021). We combine ProteomeLM embeddings with attention-derived features and achieve AUPR = 0.986, F1 = 0.923, MCC = 0.917, and AUROC = 0.997, outperforming InterSPPI (Yang et al., 2020), LSTM-PHV (Tsukiyama et al., 2021), and STEP (Madan et al., 2022) on all reported metrics. Our model and PLM-interact (Liu et al., 2025) both report near-perfect scores. Such results are inconsistent with the known difficulty of virus-human PPI prediction. We attribute them to sequence-identity leakage between the training and test splits, a problem already identified for intra-species protein-protein interactions (Bernett et al., 2024).

**Table 1.** Virus-human PPI benchmark (Tsukiyama et al., 2021; Liu et al., 2025). AUPR and F1 score are evaluated on the test set.

| METHOD | AUPR | F1 |
| --- | --- | --- |
| INTERSPPI (YANG ET AL., 2020) | 0.810 | 0.724 |
| LSTM-PHV (TSUKIYAMA ET AL., 2021) | 0.938 | 0.911 |
| STEP (MADAN ET AL., 2022) | 0.957 | 0.915 |
| PLM-INT. (LIU ET AL., 2025) | 0.999 | 0.995 |
| **PROTEOMELM-HPI** | **0.986** | **0.923** |

### 3.5. ProteomeLM-HPI performance on a strict cross-species benchmark

We developed a stricter benchmark to address the leakage issue identified above, and to test generalization to unseen pathogens. We enforce pathogen-level holdout (unseen pathogen species at test time) and sequence-level filtering at 40% identity to reduce leakage, ensuring that folds differ both in pathogen species and in sequence clusters (highly similar sequences are filtered out across splits). We use a three-way split across 10 pathogens: each fold serves as the test set once, with the remaining two folds used for training and validation respectively. We also use random cross-species negatives at 10:1 negatives-to-positives to incorporate realistic class imbalance.

Table 2 compares two baselines based on ESM-C embeddings and two ProteomeLM-based models. “Cosine” ranks candidate interactions by cosine similarity between frozen ESM-C embeddings; “ESM-LR” trains a logistic regression on concatenation, absolute difference, and element-wise product of ESM-C embeddings. “ProteomeLM-LR” uses the same pairwise feature construction on contextualized ProteomeLM embeddings followed by logistic regression; “ProteomeLM-DM” (PLM-DM) replaces logistic regression with a small downstream model (see Figure S2).

**Table 2.** Per-species AUROC on the strict cross-species benchmark. †: small dataset (less than 200 interactions).

| Species | Fold | Cosine | ESM-LR | PLM-LR | PLM-DM |
| --- | --- | --- | --- | --- | --- |
| Influenza A | A | 0.384 | 0.643 | 0.651 | **0.717** |
| HPV-16† | A | 0.640 | 0.688 | **0.731** | 0.575 |
| *M. tuberculosis*† | A | 0.577 | 0.686 | 0.655 | **0.697** |
| *C. trachomatis* | B | 0.355 | 0.710 | 0.767 | **0.819** |
| *S. enterica* | B | 0.349 | 0.730 | **0.809** | 0.804 |
| EBV | B | 0.394 | 0.596 | 0.622 | **0.700** |
| SARS-CoV-2 | C | 0.371 | 0.729 | 0.735 | **0.761** |
| HIV-1† | C | **0.612** | 0.556 | 0.526 | 0.493 |
| HSV-1† | C | 0.172 | 0.458 | 0.358 | **0.664** |
| *Y. pestis*† | C | 0.481 | **0.677** | 0.671 | 0.673 |

Cosine similarity performs poorly across the board, and is only competitive on HIV-1, where all trained models fail, likely reflecting the small dataset size (*n* = 168 positive interactions) and the atypicality of HIV-1. ESM-LR is a strong baseline, performing best on *Y. pestis* and competitively on *S. enterica* and SARS-CoV-2. ProteomeLM-LR improves over ESM-LR on several pathogens, most notably HPV-16 (0.731 vs. 0.688) and *S. enterica* (0.809 vs. 0.730), but underperforms on HSV-1 (0.358), another small dataset (*n* = 190). PLM-DM achieves the highest AUROC on 7 out of 10 pathogens, with strong gains on *C. trachomatis* (0.819), SARS-CoV-2 (0.761), and HSV-1 (0.664). These results show that a deeper downstream network built on ProteomeLM features captures relevant signals beyond what linear decision boundaries or generic ESM representations can express. This is especially true on robust datasets with more numerous pairs.

The main failures across all models regard small datasets. The five pathogens with *n* < 200 positive interactions (see Table 2) show higher variance and more frequent underperformance, highlighting a key limitation of this benchmark.

## 4. Discussion

We showed that ProteomeLM, a proteome-scale language model trained purely on intra-species masked reconstruction, encodes cross-species HPI signal. Without fine-tuning or HPI-specific supervision, it achieves AUROC = 0.786 across ten pathogen taxa. This emergent generalization can be amplified via a LoRA adaptation using asymmetric pathogen masking and blocked pathogen self-attention, yielding a model we call ProteomeLM-HPI. This fine-tuned model generalizes across ten pathogen taxa spanning viruses and bacteria, suggesting that it captures conserved principles of HPI.

ProteomeLM-HPI processes the full combined host-pathogen proteome in a single forward pass, requiring a few minutes on a single GPU. Hence, an important practical advantage is computational efficiency, compared to structure-based methods (Evans et al., 2021) and per-pair sequence classifiers (Liu et al., 2025).

We also constructed a strict cross-species HPI benchmark, motivated by the saturation of existing virus-human PPI benchmarks. By enforcing separation between pathogens and sequence-similarity filtering, our new benchmark allows to better test for generalization to unseen pathogens. The AUROCs obtained by ProteomeLM-HPI on this benchmark indicate room for further improvement, even when using supervised models. Next, we will compare ProteomeLM-HPI with prior methods on this benchmark.

This work opens several other directions. In particular, leveraging fine-tuned attention patterns to infer which human tissues a pathogen preferentially targets (tissue tropism) is a promising direction for biological interpretation.

All code and models will be made publicly available.

## Acknowledgements

This research was partly funded by the European Research Council (ERC) under the European Union’s Horizon 2020 research and innovation programme (grant agreement No. 851173, to A.-F. B.), by the Swiss National Science Foundation (SNSF, grant No. 320030 236363, to A.-F. B.), and by the Chan-Zuckerberg Initiative (CZI, to A.-F. B.).

## Appendix

## A. Evaluation dataset provenance

Table S1 lists the source publication, experimental assay, and number of positive interaction pairs for each of the ten evaluation datasets used in all benchmarks and ablations, as well as the HuggingFace virus–human benchmark. All evaluation datasets use experimentally verified binary host-pathogen interactions; negative pairs are sampled randomly during evaluation.

**Table S1.** Provenance of the ten evaluation datasets and the HuggingFace benchmark. “Assay” abbreviations: AP-MS = affinity-purification mass spectrometry; BioID = proximity-labelling mass spectrometry; Y2H = yeast two-hybrid; IntAct = IntAct database aggregate (Del Toro et al., 2022). Positive pair counts are after UniProt-ID resolution and deduplication.

| Pathogen | Source / Study | Pos. pairs |
| --- | --- | --- |
| **Viruses** |  |  |
| *SARS-CoV-2* | Zhou et al. (2023), AP-MS | 703 |
| *HIV-1* | Jaeger et al. (2012), AP-MS | 168 |
| *Influenza A* (multi-strain) | Del Toro et al. (2022), IntAct aggregate | 832 |
| *Epstein–Barr Virus* | Del Toro et al. (2022), IntAct aggregate | 1314 |
| *HPV-16* (multi-type) | Del Toro et al. (2022), IntAct aggregate | 97 |
| *HSV-1* | Del Toro et al. (2022), IntAct aggregate | 191 |
| **Bacteria** |  |  |
| *C. trachomatis* D/UW-3 | Mirrashidi et al. (2015), AP-MS | 297 |
| *M. tuberculosis* H37Rv | Penn et al. (2018), AP-MS (MiST) | 186 |
| *Y. pestis* CO92 | Dyer et al. (2010), Y2H | 196 |
| *S. Typhimurium* LT2 | Walch et al. (2021), AP-MS + BioID | 789 |
| *HuggingFace benchmark* |  |  |
| Multiple viruses | Liu et al. (2025) | — |

## B. Training hyperparameters

**Table S2.** Training hyperparameters for ProteomeLM-HPI fine-tuning. Maximum number of steps and wall-clock time differ between ablation and final models; all other hyperparameters are shared.

| Hyperparameter | Model |
| --- | --- |
| Base model | ProteomeLM-S (36M) |
| Max steps | 20 000 |
| Wall-clock time | ~1 h |
| LoRA rank $r$ | 32 |
| LoRA $\alpha$ | 128 |
| LoRA dropout | 0.1 |
| LoRA modules | $W_Q, W_K, W_V, W_{out}$ |
| Optimizer | AdamW |
| $\beta_1, \beta_2$ | 0.9, 0.999 |
| Weight decay | $5 \times 10^{-3}$ |
| Learning rate | $10^{-5}$ |
| LR schedule | Cosine |
| Warm-up steps | 500 |
| Batch size | 4 |
| Gradient clipping | 0.5 (max norm) |
| Hardware | Single NVIDIA H100 GPU |

## C. Attention for every species

**Figure S1.**
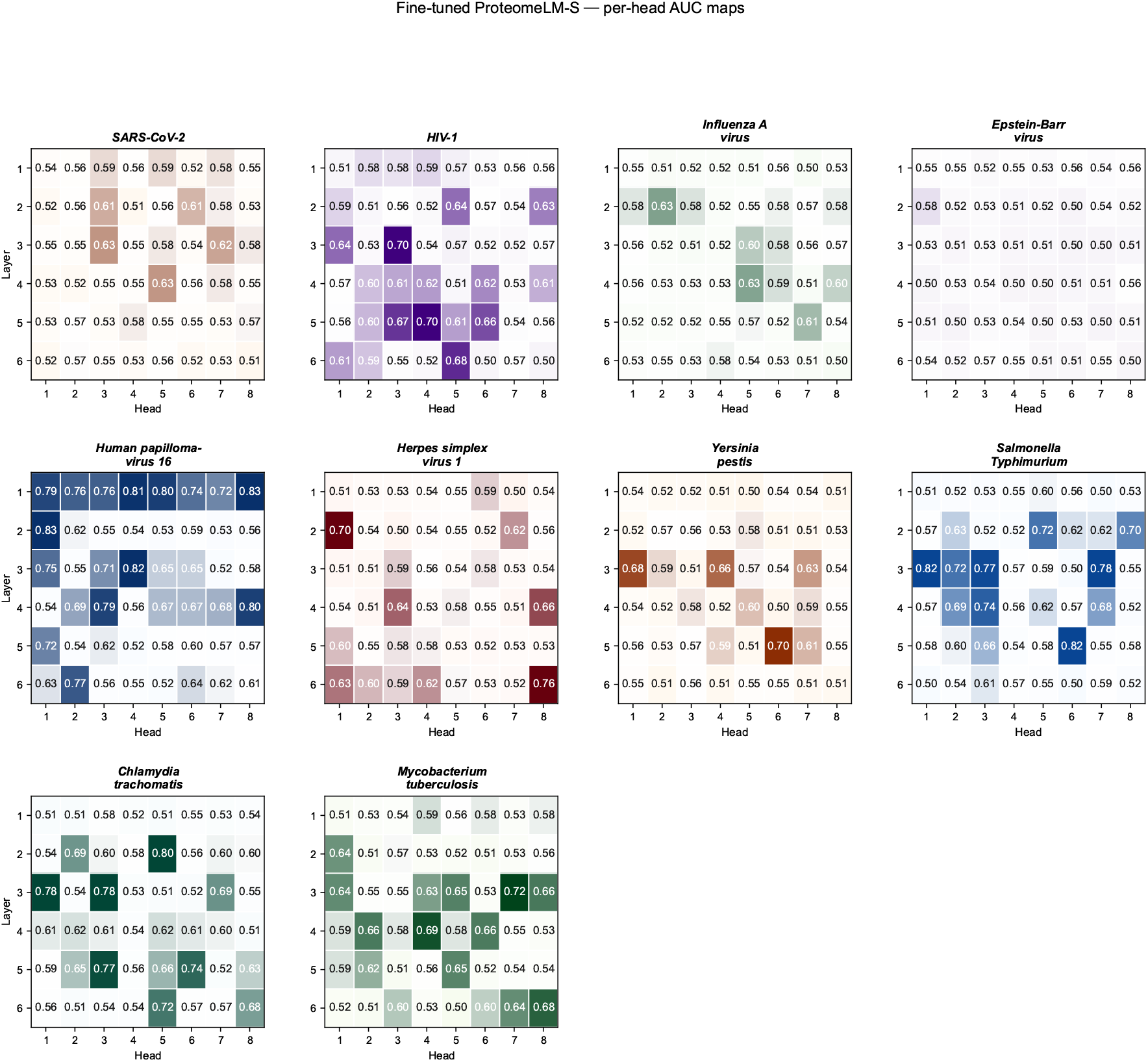
Per-head AUROC heatmaps across species for fine-tuned models. Each panel reports per-head AUROC for distinguishing true HPI pairs from random cross-species pairs in one pathogen dataset, enabling comparison of head specialization after HPI fine-tuning.

## D. Downstream model used for supervised PPI predictions

**Figure S2.**
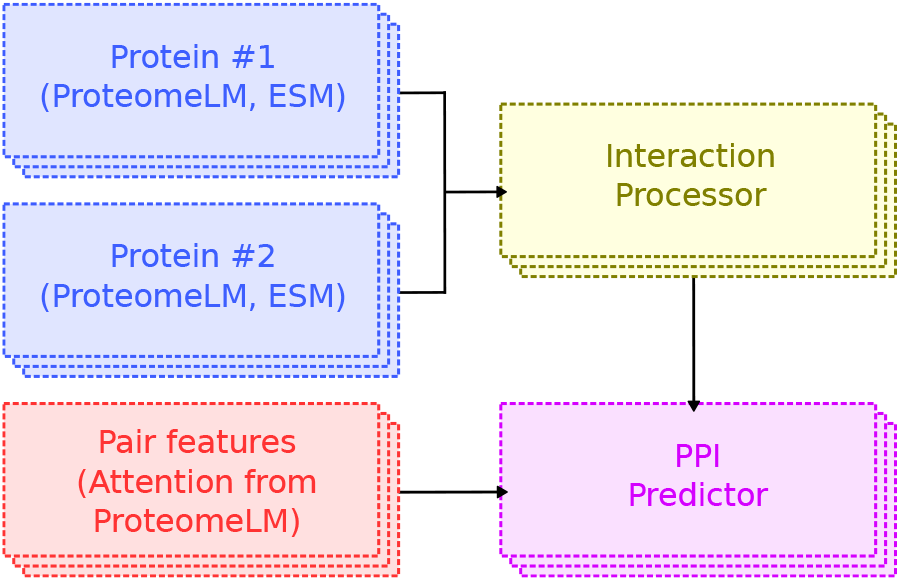
Architecture of the supervised model trained to predict protein-protein interactions (PPI). The model comprises four components. For each protein in a candidate pair, its representations from ProteomeLM-S and ESM-C are independently processed through a feature encoder.

